# SMART: An Integrated Platform Revolutionizing Cryo-Electron Microscopy Workflow from Data Acquisition to Atomic Modeling

**DOI:** 10.1101/2025.06.05.657588

**Authors:** Haibin Liu, Yangyu Wu, Wenpeng Zhang, Huaican Xu, Zhenqian Guo, Fanhao Meng, Xinyu Men, Allen Chunlong Guo

## Abstract

Recent advancements in cryo-electron microscopy (cryo-EM) have revolutionized structural biology, yet persistent workflow fragmentation between data acquisition, reconstruction, and modeling limits its full potential. This paper presents SMART, the first fully integrated cryo-EM platform unifying three critical AI-driven modules: DataSmart (automated data collection), CryoSmart (3D reconstruction), and ModelSmart (end-to-end atomic modeling). The platform embeds more than 10 distinct AI algorithms across its workflow, achieving unprecedented automation levels. These advancements not only lower the expertise barrier for cryo-EM adoption but also establish a new paradigm for AI-empowered structural biology, with broad implications for high-throughput drug discovery. This protocol provides a comprehensive delineation of the operational procedures for the SMART platform. As an example, this protocol enabled the determination of the 2.1 Å resolution structure of TRPML1, a Ca2^+^-permeable, nonselective, six-transmembrane tetrameric cation channel found in late endosomes and lysosomes (LELs) of mammalian cells. This integrated workflow enables users, including those with minimal cryo-EM experience, to efficiently complete the entire pipeline from data collection to atomic model building.

## 1. Introduction

### 1.1 Cryo-Electron Microscopy: Technological Evolution and Biological Impact

The determination of three-dimensional protein structure remains a central challenge in molecular biology, as biological function is intricately linked to the spatial arrangement of amino acid residues. Elucidating these structural configurations is imperative for deciphering enzymatic mechanisms, ligand interactions, and cellular signaling pathways at atomic resolution. This imperative has driven transformative developments in cryo-electron microscopy (Cryo-EM), which has emerged as a revolutionary structural biology tool (Nakane et al., 2020; Yip et al., 2020).

Cryo-EM single-particle analysis (SPA) serves as a powerful approach for obtaining biological insights across diverse sample types by resolving high-resolution structures of isolated complexes (Li et al., 2020). These include viruses, membrane proteins, helical assemblies, and other dynamic and heterogeneous macromolecular complexes, spanning a broad range of molecular sizes—from approximately 50kDa to tens of megadaltons.

Cryo-EM SPA has emerged as a cornerstone technique for resolving the three-dimensional (3D) architectures of biological macromolecules, encompassing three interdependent computational stages: data acquisition, 3D reconstruction, and atomic model building (Figure 1).

**Figure 1.**
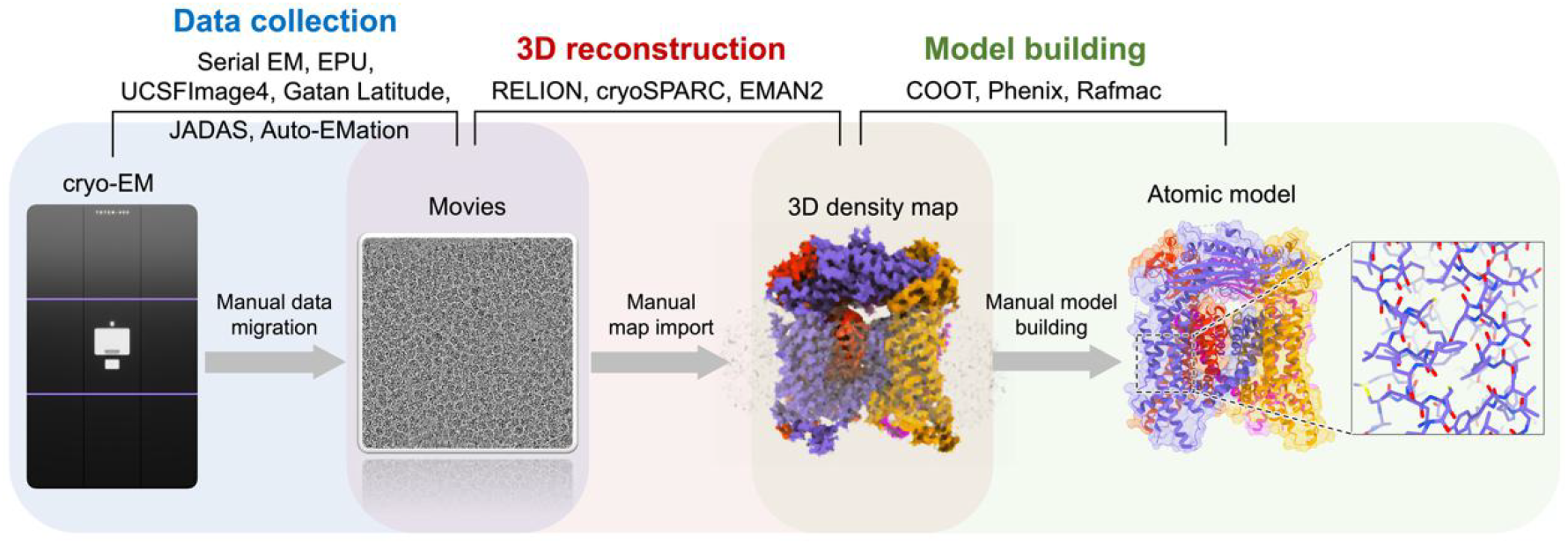
Overview of the Cryo-EM data processing using fragmented software.

During data acquisition, vitrified samples are imaged under low-dose conditions to generate micrograph movies, capturing thousands of particle projections in near-native states. These raw datasets are subsequently processed through iterative alignment and classification algorithms to reconstruct volumetric density maps. Finally, atomic models are built and refined to fit the density, elucidating molecular interactions at near-atomic resolution.

The analysis outlined above demonstrates that output of Cryo-EM merely represents the starting point of the structural pipeline. Transforming these raw data into interpretable atomic models demands a cascade of specialized computational software tools. Thus, software serves as the critical bridge transforming raw data into atomic models.

### 1.2 Fragmented Cryo-EM Software Ecosystem: Current Tools and Limitations

Modern Cryo-EM workflows rely on a patchwork of specialized software tools, each addressing discrete stages:

#### 1) Data Acquisition

While numerous Cryo-EM data acquisition software packages are publicly available—including Leginon (Cheng et al., 2021; Suloway et al., 2005), SerialEM (Mastronarde, 2003), UCSFImage4 (Li et al., 2015), TFS EPU (Deng et al., 2021; Drulyte et al., 2022), Gatan Latitude, JEOL JADAS (Zhang et al., 2009), and Auto-EMation (Lei & Frank, 2005)—these tools predominantly output raw movie data with limited post-acquisition processing capabilities. Critical workflow steps, such as parameter setting, template customization, still demand labor-intensive manual operations, often requiring too many parameters per dataset. This reliance on expert intervention introduces reproducibility challenges and workflow inefficiencies, particularly in high-throughput scenarios.

#### 2) 3D Reconstruction

Cryo-EM has witnessed remarkable advancements in 3D reconstruction software, enabling near-atomic resolution of biological macromolecules. Among the most widely used tools are RELION, cryoSPARC, and EMAN2, each offering distinct advantages and limitations. RELION employs a Bayesian approach for iterative refinement, excelling in handling heterogeneous datasets and achieving high-resolution reconstructions (Scheres, 2012). However, its computational intensity and steep learning curve often necessitate expert intervention. cryoSPARC, leveraging GPU acceleration and deep learning, significantly reduces processing time and is user-friendly, yet its performance degrades with low-SNR datasets (Punjani et al., 2017). EMAN2 provides a comprehensive suite for particle picking and 3D reconstruction but requires extensive manual tuning and lacks real-time feedback capabilities (Tang et al., 2007). While these tools excel in specific aspects of 3D reconstruction, they remain confined to singular functionalities, lacking integration with upstream data acquisition or downstream atomic modeling. For instance, RELION and cryoSPARC require manual data transfer from microscope control software, introducing inefficiencies and potential errors. Moreover, none of these tools provide real-time quality assessment during data collection or seamless transition to atomic model building, resulting in fragmented workflows that hinder throughput and reproducibility.

#### 3) Model Building

The final computational stage in cryo-EM-based structural determination is atomic model building, where molecular coordinates are iteratively fitted into reconstructed density maps to elucidate atomic-level interactions.

While conventional model-building approaches employ physics-based energy minimization algorithms—incorporating electrostatic potentials, van der Waals interactions, and covalent bond geometry constraints—they remain heavily dependent on manual intervention for initial model placement, density interpretation, and stereochemical validation (Baker et al., 2011; DiMaio et al., 2011; Lindert et al., 2012; Terwilliger et al., 2018). This reliance on expert curation not only limits throughput but also introduces subjective biases, particularly in low-resolution regimes (<4 Å) where density features are ambiguous.

To overcome these constraints, deep learning (DL)-based methodologies have emerged as transformative solutions, leveraging neural networks to automate feature extraction, model initialization, and refinement. These approaches demonstrate superior performance in handling noisy or incomplete density maps while significantly reducing manual intervention, as evidenced by recent benchmarks showing 40–60% reductions in user-dependent steps compared to conventional pipelines (Chen et al., 2016; Jamali et al., 2024; Ma et al., 2012; Si et al., 2012).

However, these methods invariably require exporting density maps from 3D reconstruction software and importing them into model-building systems. Prior to import, additional preprocessing steps—often involving third-party tools—may be necessary to optimize the density maps. This fragmented workflow necessitates repetitive data transfers and format conversions, creating inefficiencies. Moreover, if the initial model-building results are unsatisfactory, researchers must revert to the reconstruction stage for recalculation, followed by another round of model building and validation. Such iterative cycles significantly impede the throughput and operational efficiency of cryo-EM structural determination pipelines.

### 1.3 SMART: An Integrated Solution for End-to-End Cryo-EM

Despite significant advancements in Cryo-EM SPA, the lack of fully integrated computational solutions continues to impede its potential for high-throughput structural biology. Existing software packages, while excelling in automating isolated workflow components—such as data acquisition or 3D reconstruction—fail to bridge critical gaps between raw micrograph collection, density map refinement, and atomic model building. This fragmentation necessitates manual data transfer, format conversions, and redundant parameter tuning, resulting in inefficiencies that compound with dataset scale.

To address these limitations, we present SMART, the first end-to-end computational platform unifying the entire cryo-EM SPA pipeline into a cohesive, AI-driven workflow. The SMART platform integrates three core modules: DataSmart (automated data acquisition), CryoSmart (hybrid neural network-enabled 3D reconstruction), and ModelSmart (deep learning-based atomic modeling).

The platform’s three computational modules are interconnected via a standardized data stream, enabling closed-loop optimization where output from one module directly informs parameter adjustments in upstream/downstream processes. This innovation not only democratizes cryo-EM accessibility but also establishes a new paradigm for scalable, AI-powered structural determination, with transformative implications for structural biology.

## 2. Protocol

### NOTE

The SMART platform is structured into three core modules: DataSmart (automated data acquisition), CryoSmart (hybrid neural network-enabled 3D reconstruction), and ModelSmart (deep learning-based atomic modeling), with data serving as the communication interface between these software components, as illustrated in Figure 2.

**Figure 2.**
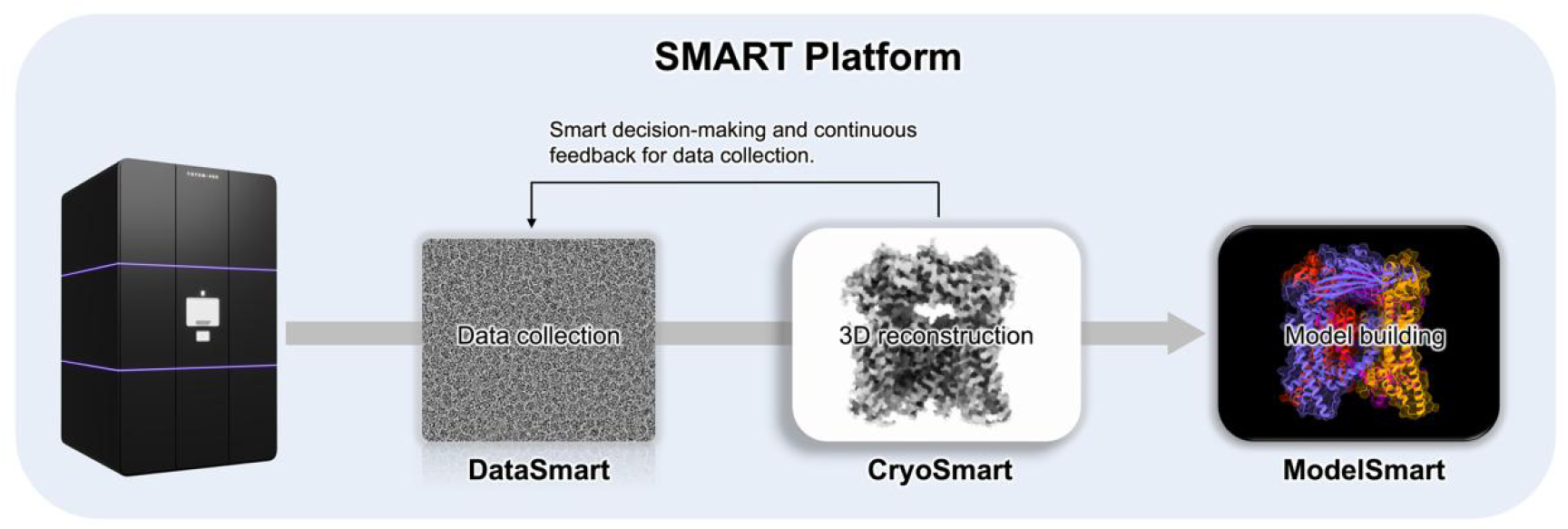
Architecture of SMART Platform for End-to-End Cryo-EM.

The SMART system adopts a Browser-Server (B/S) architecture, enabling users to perform all operations through a web interface, thereby ensuring usability and operational accessibility.

### 2.1 DataSmart (automatic data acquisition)

#### 1. Project Initialization

To begin, create a new project within the DataSmart software platform. Users may choose to inherit configuration settings from a recent project to streamline the setup process. This allows for the reuse of parameters inherited from previous projects, which can be directly applied to the electron microscope without the need for configuration from scratch.

#### 2. Beam Shift Inspection

##### Note

This adjustment is performed within the TEM control software of the electron microscope.

Beam shift must be assessed across magnifications using the Screening module. Start from low magnification and proceed to higher levels:

1. Low Magnification Check (Atlas and GridSquare): Capture images at both the Atlas and GridSquare magnifications. Verify that the images are properly illuminated and free of beam shift. Ensure that the center of the image is not positioned over the grid bars.
2. DataAcquisition Magnification Check (e.g., 60K, 100K): Acquire images at high magnifications to evaluate beam alignment. If imaging fails or significant beam shift is observed.
3. Hole Eucentric Magnification Check (typically 8K):

Acquire images at this intermediate magnification. If the beam is not centered, adjust the Beam Shift until the illumination spot is centered. Do not modify the aperture position. Ensure that the aperture size matches that used during DataAcquisition magnifications.

### 3. Z-Height Adjustment

Navigate to the AutoFunction page to perform both coarse and fine height adjustments:

1. Auto-Eucentric (Coarse Adjustment):
  1. In the Screening page, move the stage to a clean carbon film area under the Hole/EucentricHeight magnification.
  2. Open the Auto-eucentric module on the AutoFunction page and click **Start**.
  3. Monitor the log panel for feedback. If unsuccessful, consider relocating to a different area.
2. Auto-Focus (Fine Adjustment):
  1. Switch to the Autofocus module within the AutoFunction page and click **Start**.
  2. Review log messages. If autofocus fails, adjust parameters as indicated or repeat the Auto-eucentric procedure.

### 4. Magnification Alignment Check

Use the Screening page to evaluate alignment between low and high magnifications:

Alignment Inspection:

1. At the Data magnification, locate a distinguishable feature (e.g., ice contamination) and capture an image.
2. Capture an image at the Hole/EucentricHeight magnification and compare the central region of both images.
3. If misalignment is observed between Data and Hole magnifications, proceed to calibration. Otherwise, check alignment between Hole and GridSquare magnifications. If all levels are aligned, calibration is not necessary.

Calibration Procedure:

1. Navigate to the Calibration page and open the Magnification Shift module. Start from the Data magnification or the lowest misaligned level identified.
2. Capture a high-magnification image containing a distinct feature.
3. Capture a low-magnification image and assess the displacement between them. In the low-mag image, right-click the location that corresponds to the feature’s center in the high-mag image, then select **MoveHere** from the dropdown menu. Re-capture the high-magnification image to confirm alignment. Repeat if necessary. Once aligned, click **Next** to proceed to the next magnification pair.
4. Repeat Step 4 as needed (high-mag image does not need to be reacquired).

### 5. Preparation for Data Collection

This stage includes image acquisition at multiple levels and hole selection:

1. Acquire Atlas: In the Atlas module, users may either leave the field blank to capture all 81 tiles or input a number to limit the total tiles acquired.
2. Square Selection: Navigate to the Square Selection interface. Use right-click to select suitable squares.
3. Square Imaging: Proceed to the Hole Selection page and click **Prepare All**. Monitor the right-side log window for completion.
4. Hole Selection: In the image list, select the captured square image. For squares with clearly visible holes in the image, the “Find Holes” function can be directly used to perform AI-based automatic hole selection. However, in cases where the grid has excessively thick ice or significant contamination, the AI-based method may result in inaccurate hole selection. In such scenarios, the “Seed-Based Selection” function can be used to manually select holes in batches, as illustrated in Figure 3. The overall process of Seed-Based Selection is as follows:
  1. Select a central hole.
  2. Choose two adjacent holes forming a right angle.
  3. Select two distant holes along the directions defined by the second and third holes.
  4. Click **Submit** to generate a result based on the seed holes.
  5. For a different square, enable Move All in the Seed-Based Selection module. Drag one of the five seed holes to a clear hole and click **Submit** to reduce selection steps.
5. Template Definition:

**Figure 3.**
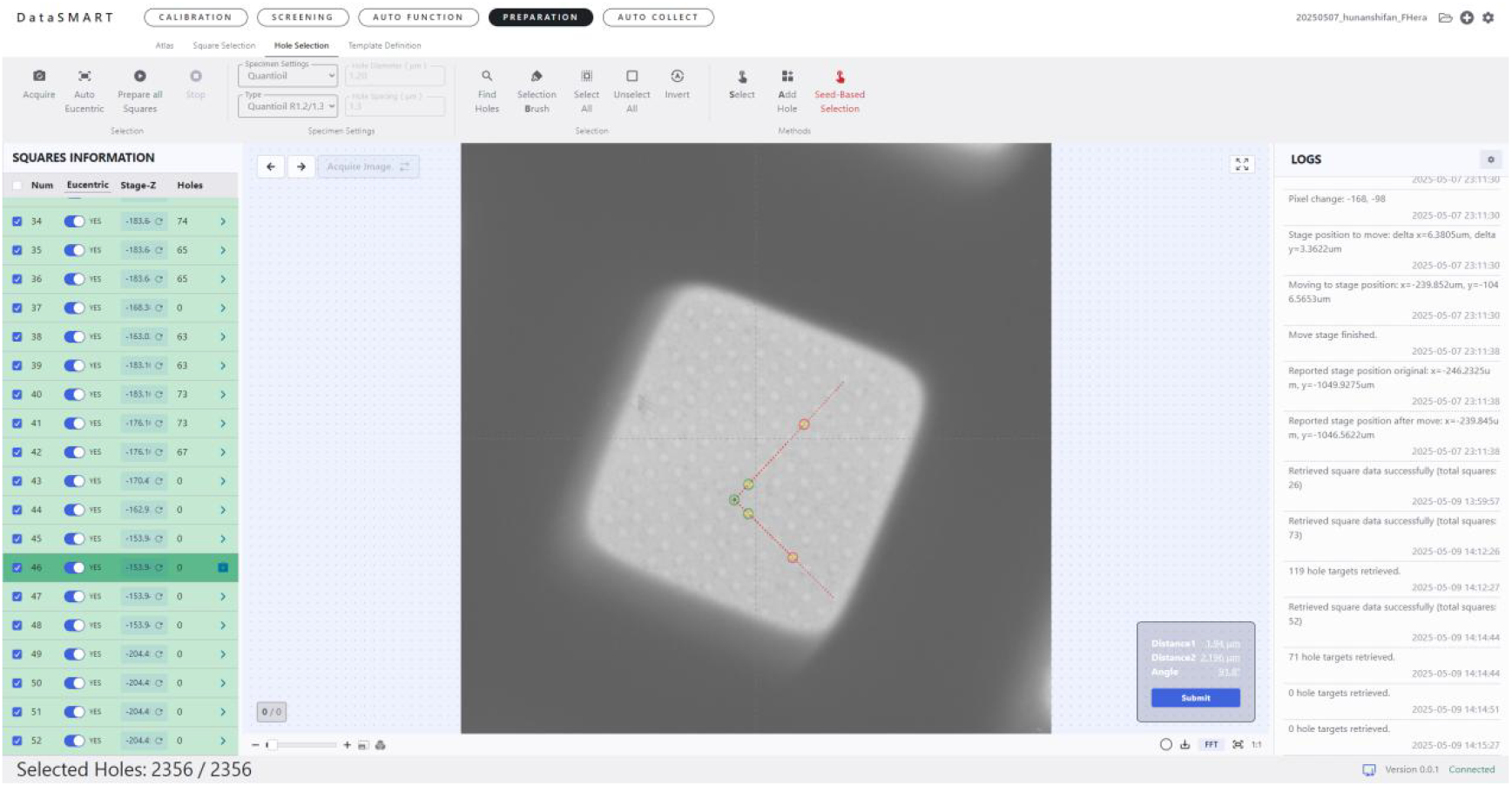
Seed-Based Selection for semi-automatic five-hole-assisted selection. The Seed-Based Selection tool is highlighted in red in the toolbar. The green part on the left side is the square list for which the image shooting has been completed. The image area in the middle shows the image corresponding to the currently selected square. On this image, five holes can be manually selected. After the selection is finished, click the blue button “**submit**”, and the number of the selected holes will be displayed in the log column on the right side.

Navigate to the **template definition** page and click **acquire** to get an image. It is essential to ensure that the current image contains a complete hole. If not, right-click on the target area and select **MoveStageTo** to reposition the stage, then reacquire the template again.

The imaging position and autofocus location can then be specified by dragging directly on the interface. Additionally, parameters such as the number of frames per hole and the desired defocus value for data collection (e.g., –1.5 µm) can be configured via the input fields, as illustrated in Figure 4:

**Figure 4.**
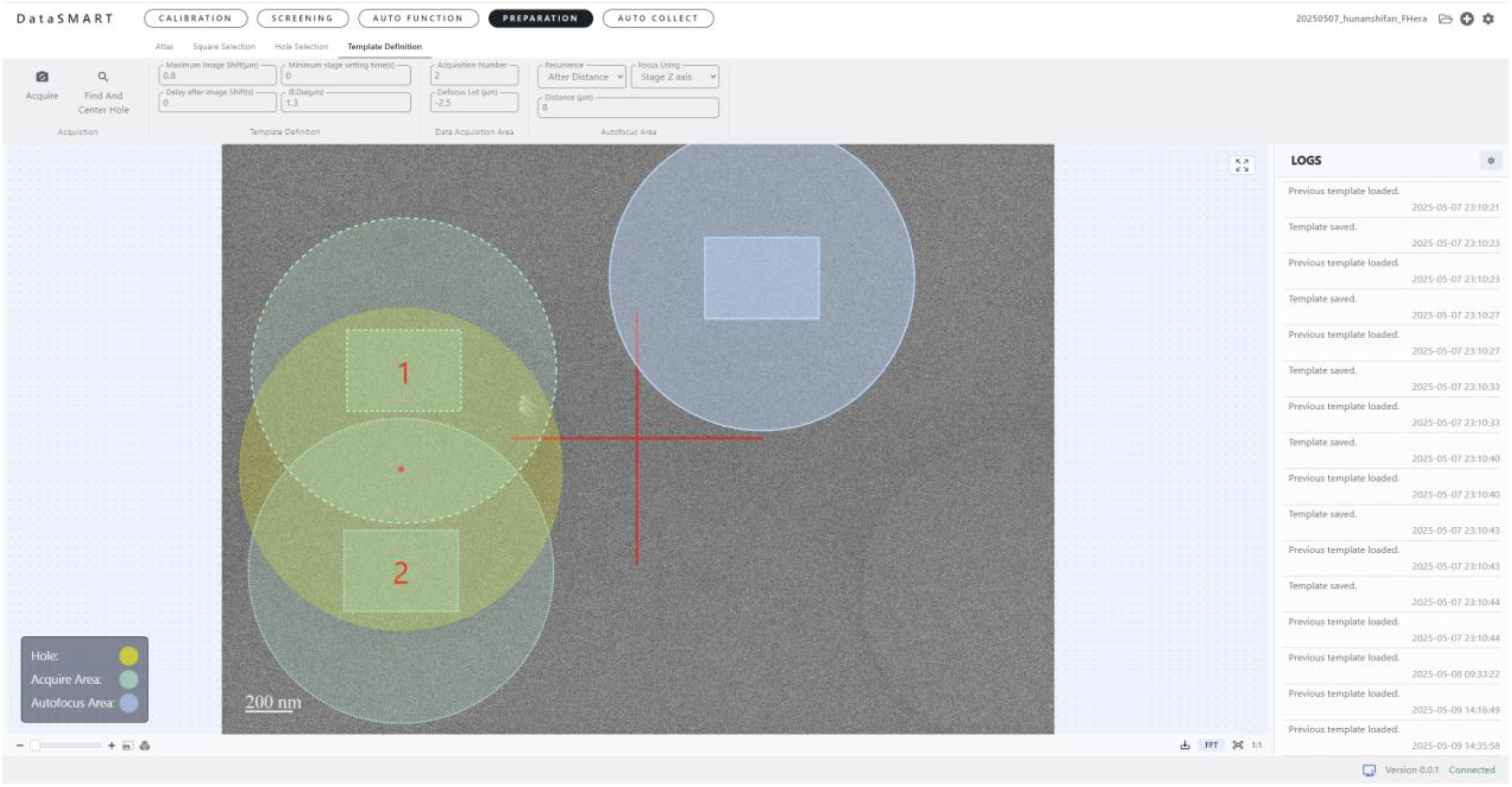
The template definition module includes an acquire button, which initiates image acquisition. In the parameter panel, users can specify the number of images to be captured within each hole, as well as the desired defocus values—either a single value or multiple values. In the image display area, three types of colored circles represent different elements: yellow for the hole, light green for the beam size, and blue for the autofocus area. The rectangle indicates the field of view of the camera during acquisition.

### 6. Beam Shift Re-Check

Return to Step 2 to reassess beam shift after all preliminary steps are completed.

### 7. Astigmatism and Coma Correction

Move the stage to a clean area of carbon film. Adjust the height via AutoFocus to the desired defocus (e.g., -1.5 µm).

Autostigmate:

Enter the Autostigmate module on the AutoFunction page and click **Start**. Monitor the right-side log for Ctffind output including astigmatism and resolution fit values. If the astigmatism remains high, repeat the process in a different area with stronger signal.

Autocoma:

If Autostigmate results remain suboptimal after multiple attempts, run Autocoma: Enter the Autocoma module and click **Start**.

### 8. Automated Data Collection

Navigate to the AutoCollect page and click **Start**. The system will automatically perform emission and energy filter calibrations (AutoEmission and AutoZLP).

Subsequently, data acquisition proceeds based on parameters defined in Screening. DataAcquisition and hole selections made in Preparation (step 5) part. Figure 5 illustrates the automatic data collection process.

**Figure 5.**
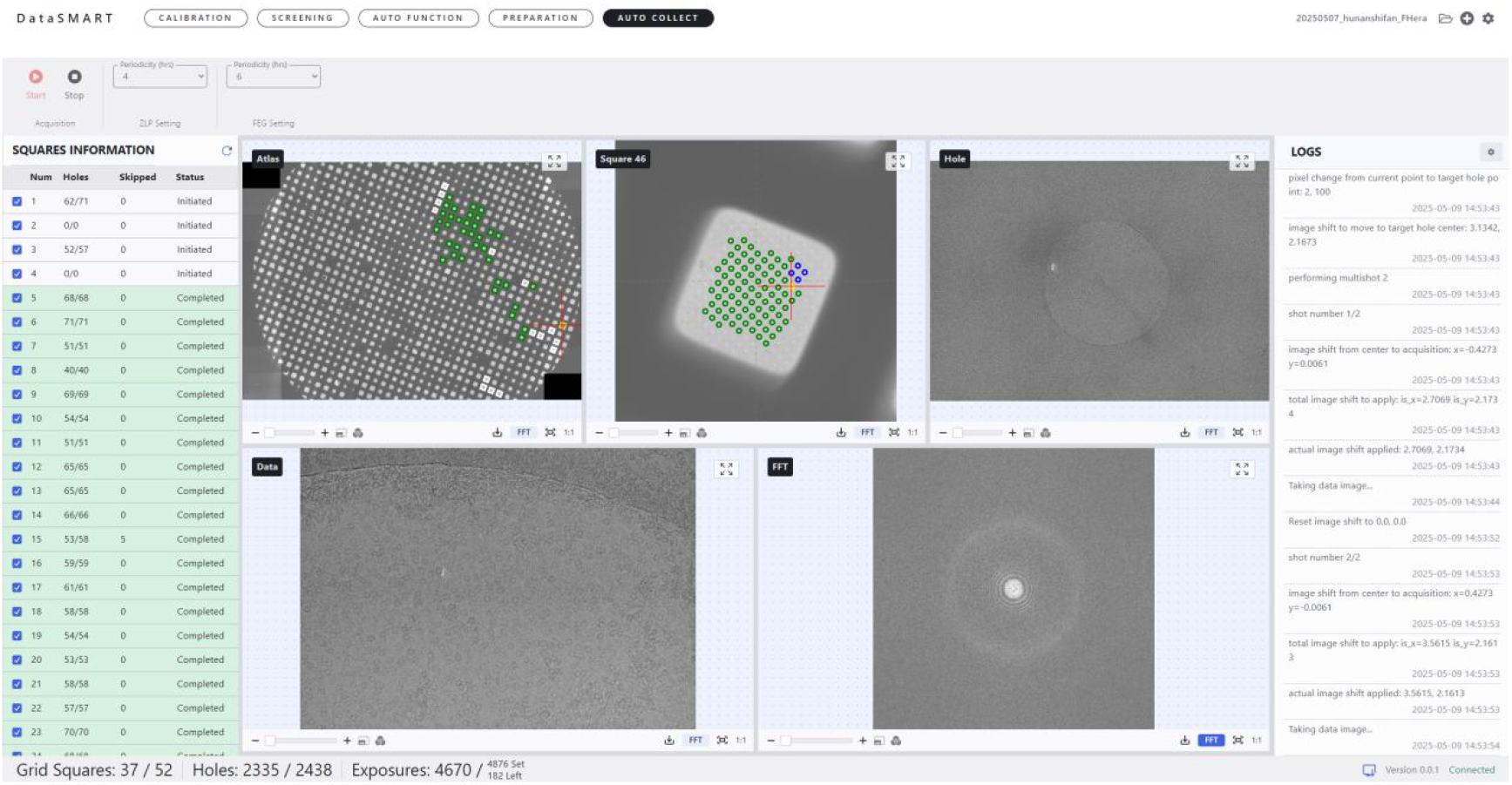
The Auto Collect module serves as the primary data acquisition module, enabling real-time monitoring of the data collection process. It provides detailed information on the specific square, group, and hole being collected, as well as the total number of images acquired. Additionally, the module displays each captured image in real time along with its corresponding Fast Fourier Transform (FFT) image.

### 9. Instant Remote Monitoring

Beyond tracking collection progress, the module also performs real-time quality assessment of the acquired data, evaluating parameters such as astigmatism and defocus. Furthermore, the system allows instant sharing of data quality information via a URL, enabling relevant users to access real-time collection updates directly from a PC or mobile device. An example of this functionality is illustrated in figure 6 below.

**Figure 6.**
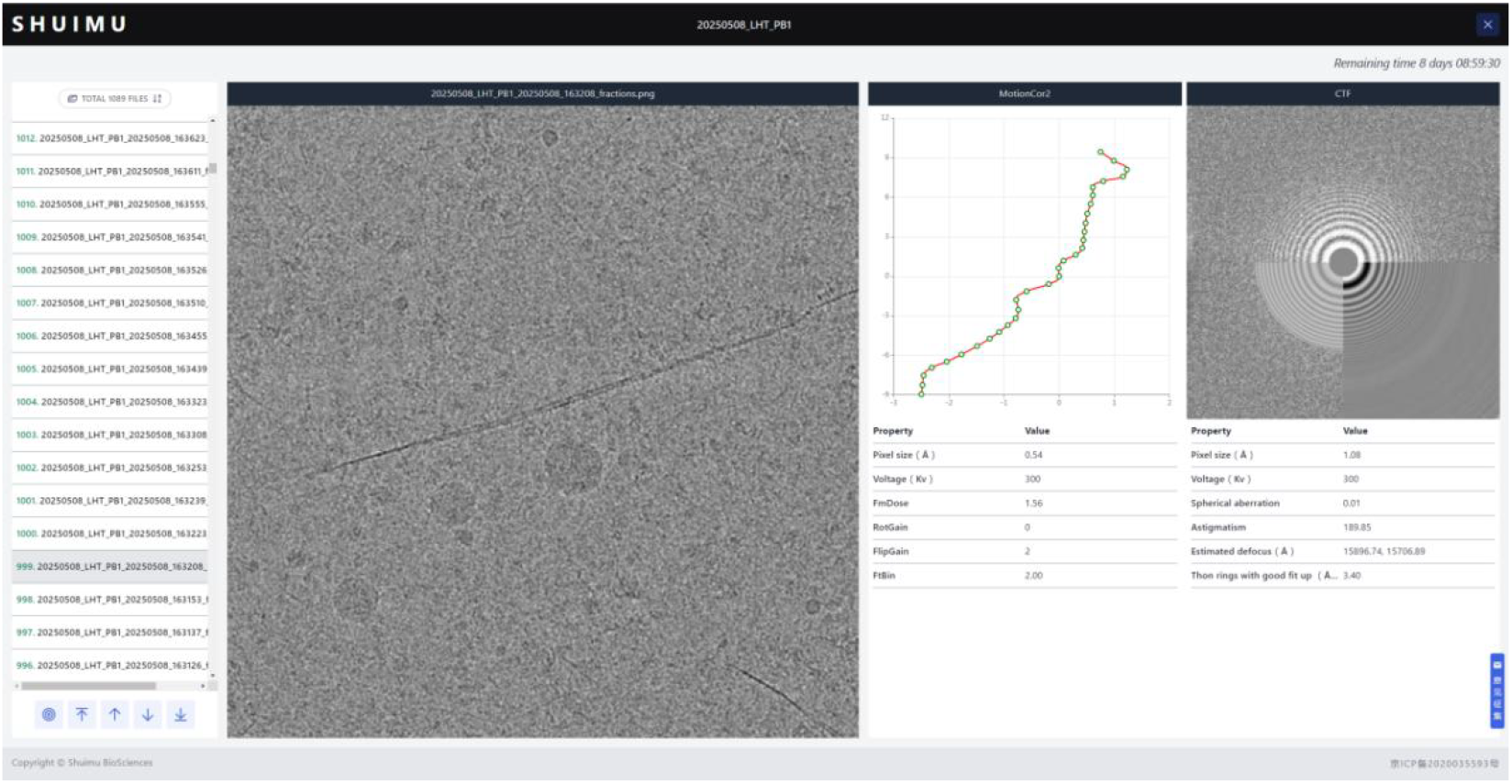
Real-Time Monitoring of Data Collection Progress and Quality in DataSmart: On the left is the list of currently collected images; The middle shows the currently selected image, and on the right side, the image motion correction and CTF information are respectively displayed.

### 2.2 CryoSmart (3D reconstruction)

CryoSMART performs three-dimensional reconstruction based on the previously acquired image datasets. The overall workflow includes project initialization, 2D micrographs import, contrast transfer function (CTF) estimation, particle picking, particle extraction, 2D classification, selection of 2D class averages, initial 3D model generation, and 3D refinement, figure 7:

**Figure 7.**
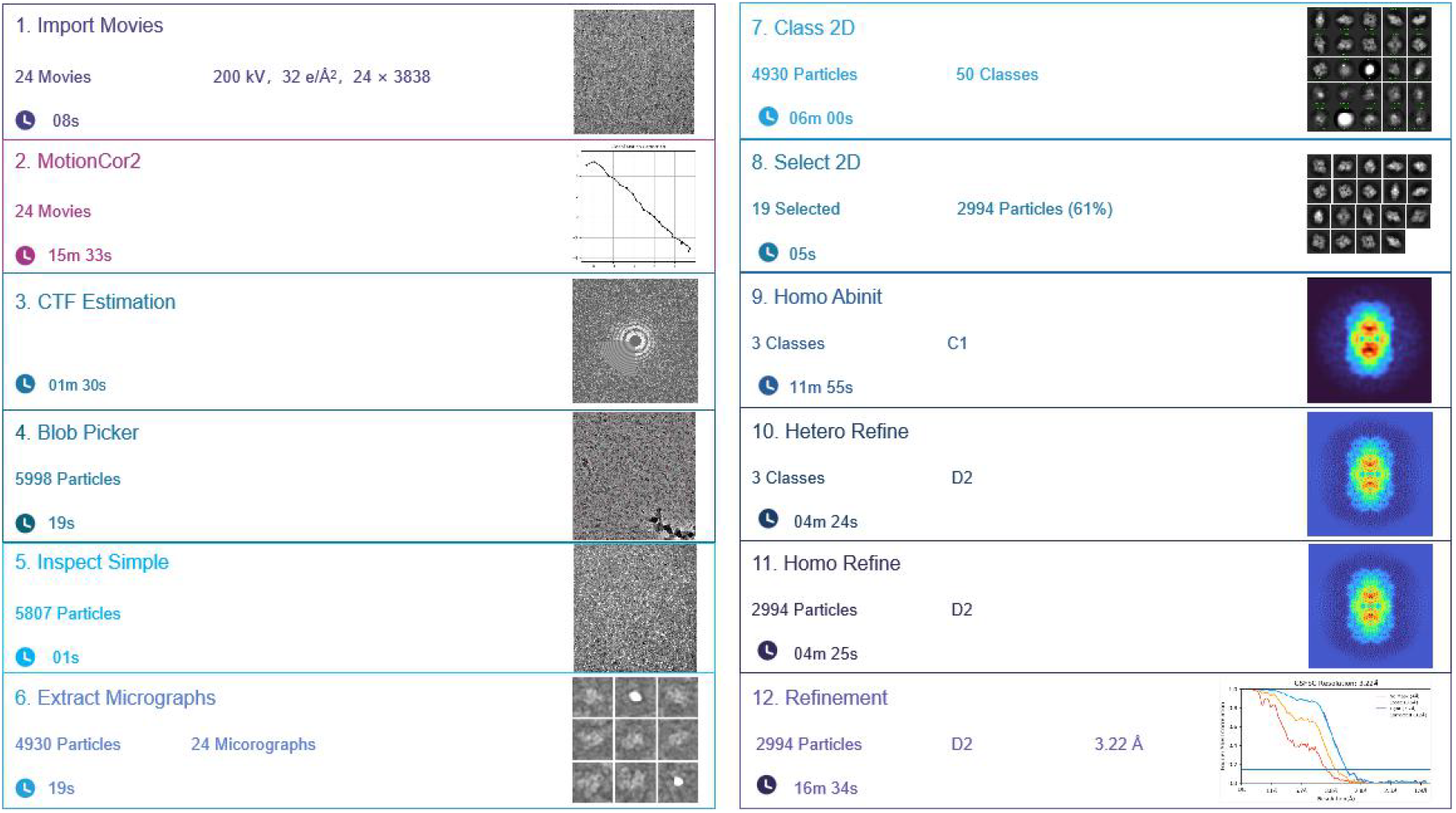
Example workflow for processing the EMPIAR-10204 dataset using CryoSmart

1. Navigate to the CryoSMART homepage and click on **Projects** to access the project management page. Click **NEW PROJECT** to create a new project.
2. A pop-up window will appear prompting the user to name the project and specify the storage location. After entering the required information, click **CREATE PROJECT** to finalize project creation.
3. Click the Open button next to the newly created project to access its workspace. Within the project page, click **NEW Experiment** to initiate a new experiment. Note: In CryoSMART, a project refers to the overall data processing session or study, while an experiment denotes a specific data processing workflow within that project. Multiple experiments can be managed under one project.
4. Next, create and execute jobs for each processing step:

1. **Importing files:** Click **New Job**, then select Import in the pop-up window. Choose the appropriate file type to be imported and click **Create** to initiate the job. A parameter input panel will appear. Import the .mrc files obtained during the data acquisition stage. Enter the required parameters such as pixel size, acceleration voltage, and the number of CPUs needed for computation; Next, click **Queue Job** to execute the job. Note:
  1. To terminate a running job, click **Kill**, then click **Clear** to reset the job status to Building.
  2. For completed or Queued jobs, simply click **Clear** to reset the job status to Building.
2. **CTF Estimation**
  1. Click **New Job**, select the CTF estimation job type in the pop-up window, and click **Create** to initiate the job.
  2. In the parameter input panel that appears, click the corresponding **Connect** button for Exposures, and select the input files in the pop-up window; then, Enter the required parameters..
  3. Finally, click **Queue Job** to run the job.
3. **Manual Picker**
  1. Click **New Job**, select Particle Picking job type and Manual Picker file type in the pop-up window, and click **Create** to initiate the job. The purpose of this step is to determine the approximate size of protein particles, which will serve as a reference for setting the particle diameter range during the subsequent *Blob Picker* step.
  2. In the parameter input panel, click the corresponding **Connect** button for *Micrographs (Exposures)*, and select the input files in the pop-up window. *Particles* input is optional at this stage.
  3. Enter the required parameters.
  4. Click **Queue Job** to run the job. An interactive interface will appear, allowing for parameter adjustments. Use the **left mouse button** to select particles and the **right mouse button** to deselect them. **Note:** View and measure particles in 3–4 micrographs (a minimum of three is recommended by the software) to determine the appropriate particle diameter range.
  5. Click **Done Picking! Extract Particles** to proceed with particle extraction and execute the job.
4. **Blob Picker**
  1. Click **New Job**, select **Particle Picking** job type and **Blob Picker** file type in the pop-up window, and click **Create** to initiate the job.
  2. In the parameter input panel, click the corresponding **Connect** button for *Micrographs (Exposures)* and select the input files in the pop-up window.
  3. Enter the required parameters, then click **Queue Job** to run the job.
5. **Inspect Picks**
  1. Click **New Job**, select **Particle Picking** job type and **Inspect Particle Picks** file type in the pop-up window, and click **Create**.
  2. In the parameter input panel, click the **Connect** buttons for both *Micrographs (Exposures)* and *Particles*, then select the corresponding input files in the pop-up windows.
  3. Enter the required parameters and click **Queue Job** to execute the task.
  4. After execution, an interactive interface will appear for parameter adjustments. Examine 3–4 images to ensure correct particle selection. Then, click **Done Picking! Output Locations** to proceed.
6. **Extract from Micrographs**
Purpose: To extract individual particles from the micrographs.
  1. Click **New Job**, select **Particle Picking** job type and **Extract From Micrographs** file type in the pop-up window, and click **Create**.
  2. In the parameter input panel, click the **Connect** buttons for *Micrographs (Exposures)* and *Particles*, and select the corresponding files.
  3. Enter the required parameters and click **Queue Job** to run the job. After importing the images, fill in the necessary parameters. Set the Extraction box size (pix) to 2–3 times the particle diameter. The Fourier crop to box size should be set to 1/4 to 1/2 of the extraction box size. Note: Reducing the Fourier crop size helps minimize the computational load on the GPU.
7. **2D Classification** Purpose: To classify particles based on their appearance in 2D images. Click **New Job**, select **2D Classification** file type in the pop-up window, and click **Create**. In the parameter input panel, click the Connect button for Particles and import the relevant files. Enter the required parameters and click **Queue Job** to execute the classification. Note: It is recommended to select 50–200 classes. For high-quality datasets with fewer particles, use fewer classes; for high-quality, large datasets, consider selecting 150–200.
8. **Select 2D Classes** Purpose: To select high-quality 2D class averages. Click New Job, select **Select 2D classes** in the pop-up window, and click **Create**. In the parameter input panel, click the **Connect** buttons for Particles and Templates, and import the required files. click **Queue** Job. After approximately 3 seconds, the job will enter a Waiting state. Click **Waiting** to begin selection. In the displayed 2D classification results, manually select clear and well-defined classes. Click **Done** to proceed.
9. **Initial Model Generation** Create new job. select **Abinit Reconstruction** in the pop-up window. In the parameter input panel, click the **Connect** button for Particles and import the selected particle file from the previous step. Enter the number of classes to be initialized as well as the symmetry parameter. Click **Queue Job** to run the job.
10. **Heterogeneous Refinement**
Note: Choose between Homogeneous Refinement and Heterogeneous Refinement depending on whether the protein sample consists of a homogeneous or heterogeneous complex.
  1. Click **New Job** to create a Heterogeneous Refinement job. In the parameter input panel, click the **Connect** buttons for Particles and Volume, and select the input files.
  2. Enter parameters. In Heterogeneous Refinement, the Symmetry setting should reflect the actual symmetry of the protein (e.g., C1). Click **Queue Job** to begin refinement.
11. **Homogeneous Refinement**
  1. Click **New Job**, select the appropriate job type, and create a Homogeneous Refinement job. In the parameter input panel, click the **Connect** buttons for Particles (required), Volume (required), and Mask (optional), then select the corresponding input files, as shown in Figure 8.
  2. Import the single-particle dataset and the best-performing volume_class model from the previous Heterogeneous Refinement step.
  3. Enter the required parameters. As in the previous step, the Symmetry setting should be specified according to the actual properties of the protein (e.g., O or C1). Click **Queue Job** to start the refinement.
  4. After the job is completed, click on “**Output**” to directly preview the calculated density map in real-time on the browser page, as illustrated in Figure 8.
12. **Map Enhancer**
Note: Map Enhancer is designed to generate post-processed maps from raw input volumes. It performs optimally when using half-maps as input. The specific model architecture used by SMART is DeepEMhancer (Sanchez-Garcia et al., 2021), a deep learning-based method trained on paired datasets of experimental cryo-EM volumes and their atomic model-refined counterparts. Please note that post-translational modifications and ligand molecules were excluded from the training data, and thus the model may not accurately enhance such features. Essentially, Map Enhancer applies non-linear enhancements to cryo-EM maps, delivering two primary outcomes:
  1. Local sharpening similar to conventional post-processing methods.
  2. Automated masking and denoising of the input cryo-EM maps.
    1. Click **New Job**, select the appropriate job type, and create a Map Enhancer job.
    2. In the parameter input panel, import the Volume and Mask data from the previous Homogeneous Refinement step into the corresponding DeepEMhancer input fields.

**Figure 8.**
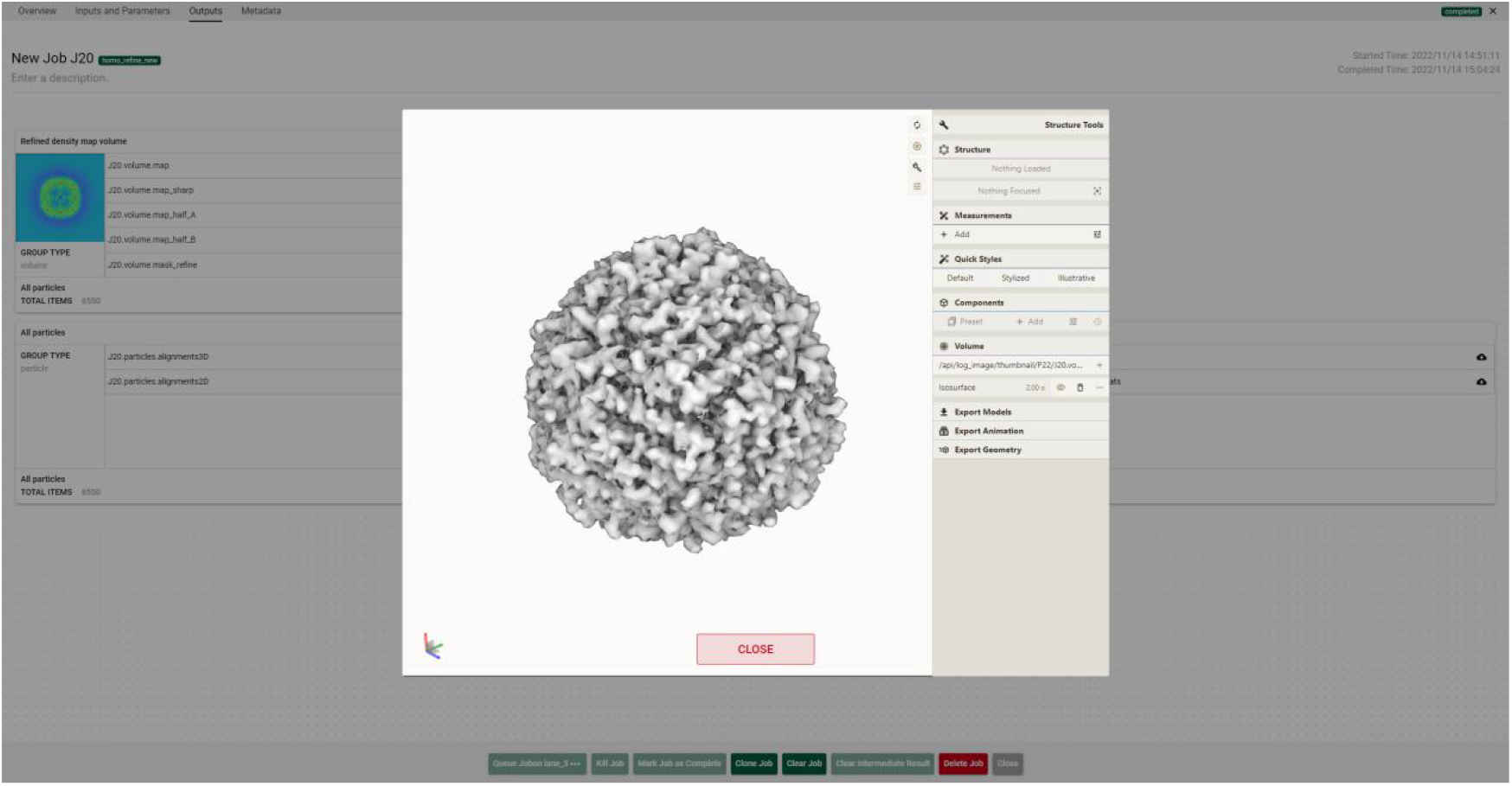
3D density map of Apoferritin viewed directly within CryoSmart. The computed density map display enables operations such as rotation and stretching of the density map via mouse interaction.

Alternatively, you can use the Import Files → Map Data Path option to select files.

**ModelSmart (Atomic Modeling)**

Building upon our previously developed SMARTFold (Li et al., 2023) framework, the ModelSmart system represents a specialized advancement in atomic modeling for cryo-EM, extending its capabilities to diverse biomolecular targets, including proteins and nucleic acids (RNA/DNA).

After logging into the system, users can upload the density map on the left panel and input the required parameters to submit a task for ModelSMART processing.

The required input parameters are as follows:

**Cryo-EM Density Map**: Upload the density map file in .mrc, .ccp4, or .map format (file size must be less than 100 MB).

Note:

To ensure optimal model performance and minimize the risk of errors, it is strongly recommended to inspect the density map before uploading.

1. Density Map Cropping: Cropping the density map helps remove redundant regions, thereby significantly reducing the file size and preventing potential memory overflow during model processing.
2. Mirror Inversion of Structures: Cryo-EM density maps may sometimes be mirrored. If a mirrored structure is detected, it must be flipped accordingly.

To determine whether a map is mirrored, examine the α-helices in the protein structure—under normal conditions, they should exhibit a right-handed configuration, spiraling upward in a counterclockwise direction.

**Sequence File**: Upload the corresponding sequence file, or manually input the sequence if proteins, nucleotides, or heteromeric/homomeric forms if involved.

**Protein Name** *(optional)*: Enter the name of the protein. **Resolution**: Specify the exact resolution of the density map. **Partition**: Select the GPU computing node for job execution.

**Number of CPUs**: Enter the number of CPUs to be used (maximum of 8). Finally, click the **SUBMIT** button to initiate the job.

After the job is submitted, the predicted atomic model is rendered in an interactive 3D visualization environment. Users can freely rotate, zoom, and manipulate the model in real time, enabling comprehensive inspection of structural features from multiple perspectives. This functionality facilitates an intuitive evaluation of model quality, spatial conformation, and atomic-level detail, supporting downstream analysis and validation of the predicted structure, as illustrated in the figure 9.

**Figure 9.**
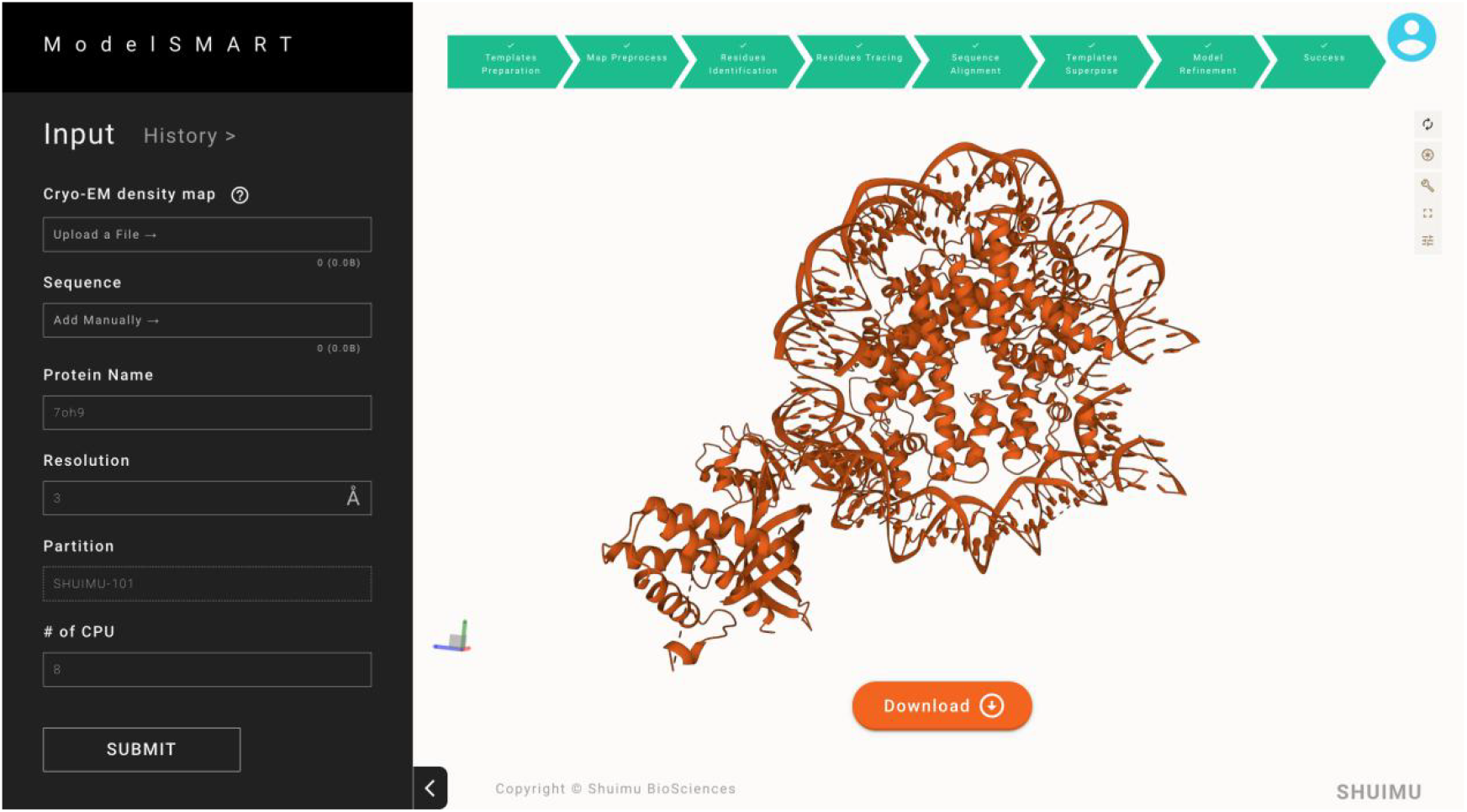
The atomic model displayed directly in ModelSmart. Through the mouse, the atomic model can be rotated and stretched, specific amino acid information can be selected, and clicking the “**Download**” button allows direct downloading of the atomic model’s PDB results.

## Representative Results

TRPML1, also known as mucolipin-1 (MCOLN1), is a lysosomal cation channel essential for maintaining ionic homeostasis, regulating vesicular trafficking, and supporting autophagy in mammalian cells (Wang et al., 2014; Xu & Ren, 2015). Dysfunction of TRPML1 due to genetic mutations has been directly linked to mucolipidosis type IV, a severe neurodegenerative lysosomal storage disorder characterized by abnormal neurodevelopment, retinal degeneration, and iron-deficiency anemia (Bargal et al., 2000; Bassi et al., 2000; Nilius et al., 2007; Sun et al., 2000). Understanding the structural basis of TRPML1 function is thus critical to elucidating its physiological and pathological roles.

A total of 1,034 movie stacks were collected at a nominal magnification of 96,000×, corresponding to a super-resolution pixel size of 0.808 Å (Figure 10A). Movies were recorded over 5.35 seconds with 32 frames per stack and an accumulated dose of 50 e− /Å^2^. Defocus values ranged from –1.0 to –1.5μm. A multi-shot acquisition strategy employing image-shift patterns was used to increase throughput and maximize microscope usage. Notably, **all data acquisition steps were fully automated using the in-house developed *SMART* platform**, which integrates hardware control with real-time image quality assessment and beam alignment correction, significantly reducing human intervention.

**Figure 10.**
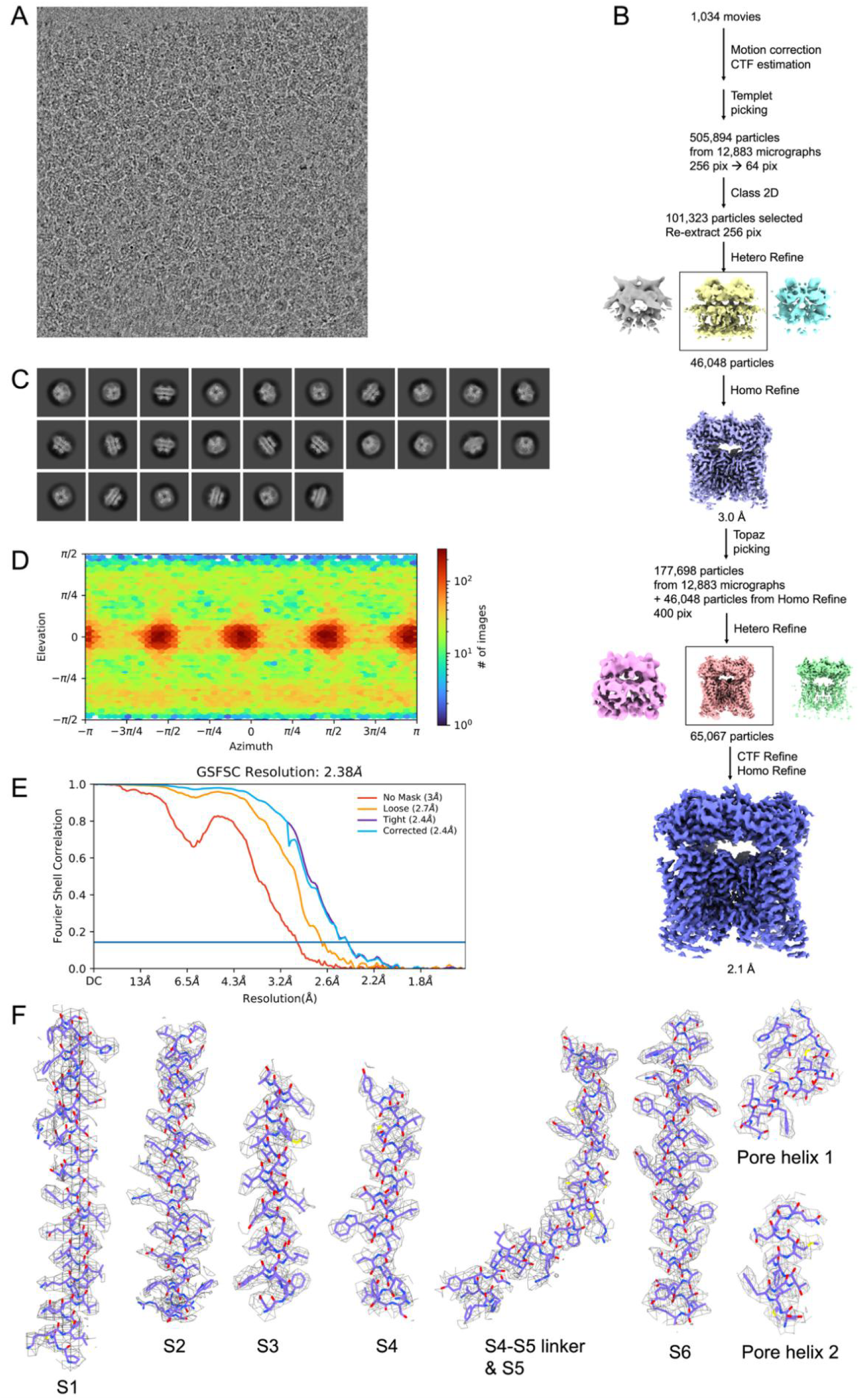
Image processing and reconstruction of human TRPML1. (A) Representative micrograph. (B) Cryo-EM data processing workflow. (C) Representative 2D classes. (D) Particle distribution plot. (E) Gold standard FSC resolution estimation. (F) Cryo-EM density map and model for alpha helices in human TRPML1 structure.

Cryo-EM image processing was carried out using our custom software suite C***ryoS***mart, which features a fully integrated pipeline for motion correction, CTF estimation, particle picking, classification, and refinement. Unless otherwise specified, all processing steps were executed within CryoSmart. An initial particle set was generated using a template-based picker seeded by 2D projections derived from a 3.72 Å reference map of human TRPML1 (EMDB-8840). After 2D classification and extensive cleaning using 4× downsampled particles, ∼101,000 high-quality particles were retained and re-extracted at full resolution for subsequent refinement. Three major classes emerged from heterogeneous refinement, with the best-resolved class (46,048 particles) subjected to homogeneous refinement, incorporating per-particle CTF refinement, aberration correction, and anisotropic magnification calibration. This yielded a reconstruction at **3.05 Å global resolution**.

To further improve the completeness and uniformity of particle picking, a **Topaz model was trained and executed directly within** C**ryoS**mart, leveraging its plug-in compatibility. This deep-learning-based model was trained on particle coordinates from the 3.05 Å structure, assuming an average of 100 particles per micrograph. The retrained Topaz model was applied to the entire dataset, yielding 177,698 particles. These, along with the previously selected 3.05 Å stack, were extracted into 400-pixel boxes and re-classified. A high-resolution subset (class 2) was identified, with duplicates removed. Final homogeneous refinement using 83,861 particles resulted in a **2.10 Å global resolution** reconstruction, enabling visualization of side-chain rotamers and detailed intra-subunit interactions of TRPML1 (Figure 10B-F).

Our results demonstrate that **SMART** platform provides an efficient, streamlined, and automated platform for high-throughput cryo-EM structure determination. The ability to complete the entire workflow—from data acquisition to near-atomic structure—within a significantly shortened timespan highlights the potential of this software ecosystem in accelerating structural studies, especially for membrane proteins with challenging behavior. By minimizing manual intervention and maximizing automation and reproducibility, our system offers a scalable solution for both academic and industrial cryo-EM applications.

## Discussion

The structural determination of biological macromolecules via cryo-electron microscopy (cryo-EM) inherently involves a protracted, multi-stage workflow requiring tight integration of computational tools and reliance on expert intervention. The SMART platform addresses this challenge by unifying data acquisition, 3D reconstruction, and atomic modeling into a single, vertically integrated framework—a critical advancement in an era of high-throughput structural biology. By embedding AI-driven algorithms across the pipeline (e.g. geometric deep learning for density-guided model building), SMART reduces manual interventions compared to conventional fragmented workflows. This automation not only lowers the expertise barrier but also enhances reproducibility. In the current landscape where cryo-EM throughput demands are rapidly rising—from academic labs to pharmaceutical pipelines—such an end-to-end solution represents a timely and critical advancement.

SMART’s server-based deployment eliminates reliance on external networks, a pivotal feature for institutions handling sensitive biomedical data. A notable advantage of SMART is its real-time image display capability, which is crucial for decision-making during data processing. While many comparable systems suffer from latency or image stuttering, SMART circumvents this by deploying a dedicated image-processing subprocess, ensuring seamless visualization. Furthermore, the platform provides a robust set of APIs, facilitating easy integration with third-party software and enabling customized extensions. This design enhances both the usability and scalability of the system for diverse cryo-EM research needs.

The successful determination of the human TRPML1 structure at 2.38 Å using SMART illustrates this potential. Importantly, the automated workflow not only accelerated the timeline from micrograph acquisition to atomic model, but also ensured consistent data quality through built-in real-time monitoring and particle picking optimization. The integration of deep learning modules, such as Topaz for particle selection and ModelSmart for atomic modeling, exemplifies how AI-driven components can transform previously manual bottlenecks into scalable, automated steps.

Yet, while SMART substantially advances cryo-EM workflow integration, several frontiers remain open for innovation. Current implementations lack closed-loop integration between data acquisition and atomic model validation. Future developments will focus on implementing real-time atomic model building during microscope operation, where AI-driven quality assessment of preliminary models (e.g., resolution estimates) dynamically determines whether sufficient data has been collected. This capability will enable automated termination of imaging sessions once predefined resolution thresholds (e.g., 3.0 Å) are statistically validated, while simultaneously optimizing beam parameters (defocus, dose rate) to maximize model accuracy. Such functionality would permit intelligent termination of data collection once predefined resolution thresholds (e.g., 3.5 Å) are statistically assured, potentially reducing beam time by 30–50%.

Moreover, as AI tools like AlphaFold (Jumper et al., 2021) and RoseTTAFold (Baek et al., 2021) reach maturity, their integration within cryo-EM pipelines opens exciting prospects. For instance, leveraging sequence-based predictions to guide initial model placement or to resolve flexible and poorly ordered regions in intermediate resolution maps (<4 Å) remains an underexploited avenue. In this context, the modular design of SMART provides a fertile ground for incorporating structure prediction modules and multi-state classification tools, enabling users to explore conformational heterogeneity with greater fidelity. Additionally, the accumulation of user-generated training data across diverse biological targets will enable the development of context-aware AI models, enhancing performance on rare or heterogeneous complexes. With the development and maturation of large language models(LLMs) like ChatGPT (Achiam et al., 2023),The integration of LLMs could further revolutionize user interaction, enabling natural language commands (e.g., “Optimize grid screening for membrane proteins”) or voice-guided workflow customization—a paradigm shift toward democratized, conversational structural biology. Finally, extending SMART’s architecture to support cryo-electron tomography (cryo-ET) workflows and multi-modal data fusion (e.g., correlative light-electron microscopy) will broaden its applicability in cellular and subcellular contexts.

In conclusion, SMART represents a foundational step toward the next generation of cryo-EM platforms: fully automated, AI-driven, and dynamically adaptive. Its deployment in both academic and industrial settings hold the promise of accelerating structural discovery pipelines, particularly for challenging targets such as membrane proteins and transient complexes. By continuing to evolve toward closed-loop control, AI-guided decision-making, and multi-modal data fusion, SMART has the potential to redefine throughput and accessibility standards in structural biology.

## Acknowledgments

We thank Dr. Jing Li from Cryo-EM Center Shuimu BioSciences for his advice for the representative dataset. We thank Dr. Yujie Liu from Protein and Crystallography Department Shuimu BioSciences for providing the hTRPML1 protein sample. We gratefully acknowledge Associate Professor Qiangfeng Zhang, Dr. Kui Xu, Associate Professor Xueming Li, and Engineer Bo Shen from the School of Life Sciences, Tsinghua University, as well as Professor Xinzheng Zhang and Dr. Chunling Wu from the Institute of Biophysics, Chinese Academy of Sciences, for their valuable support and guidance during the development of the SMART platform.

